# Fast functional annotation of metagenomic shotgun data by DNA alignment to a microbial gene catalog

**DOI:** 10.1101/120402

**Authors:** Stuart M. Brown, Yuhan Hao, Hao Chen, Bobby P. Laungani, Thahmina A. Ali, Changsu Dong, Carlos Lijeron, Baekdoo Kim, Konstantinos Krampis, Zhiheng Pei

## Abstract

**Background:** Metagenomic shotgun sequencing is becoming increasingly popular to study microbes associated with the human body and in environmental samples. A key goal of shotgun metagenomic sequencing is to identify gene functions and metabolic pathways that differ between samples or conditions. However, current methods to identify function in the large number of reads in a high-throughput sequence data file rely on the computationally intensive and low stringency approach of mapping each read to a generic database of proteins or reference microbial genomes.

**Results:** We have developed an alternative analysis approach for shotgun metagenomic sequence data utilizing Bowtie2 DNA-DNA alignment of the reads to a database of well annotated genes compiled from human microbiome data. This method is rapid, and provides high stringency matches (>90% DNA sequence identity) of shotgun metagenomics reads to genes with annotated functions. We demonstrate the use of this method with synthetic data, Human Microbiome Project shotgun metagenomic data sets, and data from a study of liver disease. Differentially abundant KEGG gene functions can be detected in these experiments.

**Conclusions:** Functional annotation of metagenomic shotgun sequence reads can be accomplished by rapid DNA-DNA matching to a custom database of microbial sequences using the Bowtie2 sequence alignment tool. This method can be used for a variety of microbiome studies and allows functional analysis which is otherwise computationally demanding. This rapid annotation method is freely available as a Galaxy workflow within a Docker image.

## Background

The collection of large scale metagenomic shotgun DNA sequence (MGS) data sets from microbial communities associated with the environment, the human body (microbiome), or from other animals has become common. The initial focus of metagenomics studies, such as the Human Microbiome Project [1] was to survey the microbial species present in various sites on and in the human body, but the focus of research has now shifted to understanding the functional role these microbes play in metabolic and disease processes. Measurement of the taxonomic composition of metagenome samples by PCR and amplicon sequencing of the 16S rDNA marker gene is inexpensive, but it is subject to bias and lacks sensitivity below the species level. Individual bacterial isolates with identical 16S genes may differ by as much as 15-30% in their genomes [2, 3], which may include toxin production, antimicrobial, or metabolic genes. Shotgun sequencing of all DNA present in a biological sample can be used for computational prediction of gene functions of sequenced DNA fragments to infer differences in the metabolic capacity of microbial communities [4].

Existing bioinformatics tools to characterize MGS data are problematic due to the large computational task of comparing millions of short reads (50 to 200 nucleotides in length) to various databases of known proteins, conserved protein motifs, or annotated complete genomes. These databases typically lack many of the gene/protein sequences from the actual microbial species present in microbiome samples, which contain organisms that cannot be cultured. BLAST [5], is the most commonly used (and the most sensitive) method to compare DNA sequences to a database but it requires hundreds of CPU hours to analyze a typical MGS sample FASTQ data file containing hundreds of millions of reads. Approaches to overcome this computational bottleneck have attempted to reduce the query data in each data file by de-duplication, or by *de novo* assembly. However, these data reduction methods themselves require substantial computational effort and can introduce bias. Other methods use faster, but less sensitive sequence matching algorithms such as BLAT [6] or RAPSearch [7] and reduced databases for functional protein identification, providing a less precise assay for microbial protein function. The popular MG-RAST webserver implements a MGS pipeline that combines aspects from all of these approaches [8], but it suffers from its own bottlenecks, since raw data must be uploaded over the internet for processing, and it has a queue which can take several weeks.

## Methods

We have created a computationally efficient pipeline for MGS analysis (MGS-Fast), which combines data cleaning, removal of human sequences, profiling of taxonomic composition, and functional profiling of microbial sequence fragments. The pipeline utilizes existing tools for quality control, sequence trimming, taxonomy, and DNA sequence alignment; but applies them in a novel manner, using rapid stringent DNA-DNA matching to previously annotated microbiome sequences to assign functions to MGS reads. We have evaluated this annotation method on existing public metagenomic data sets, simulated data from microbial and human genomes, and shotgun metagenomic data from the human liver disease study of Qin et al [8].

### Datasets

Metagenomics datasets tested on the MGS-Fast pipeline were downloaded as Illumina FASTQ files from the NCBI SRA for human skin (SRR1646957), oral (SRR769511), gut (SRR2822459), and two synthetic bacterial communities (SRR3732372 and SRR172902). Human esophagus and oral samples (dbGaP Study Accession: phs000260.v3.p2). Liver cirrhosis and healthy controls were obtained from European Nucleotide Archive Study PRJEB6337.

Additional data sets were downloaded from MG-RAST [9] for mouse gut (MG-RAST 4535626.3) and for copper mine waste (MG-RAST 4664533.3). Simulated FASTQ data files were created from the human GRCh38 reference genome and the *E. coli* K12 reference genome (GenBank: accession U00096.3) by MetaSim [10] and a random DNA sequence was created in FASTQ format by XS simulator [11].

### Data Analysis Pipeline

The MGS-Fast annotation method relies on a large database of metagenomic DNA sequences built from the Integrated Gene Catalog (IGC) of the human gut microbiome [12] and the Human Oral Microbiome Database (HOMD) [13]. This database contains *de novo* assemblies and annotations of almost 10 million “genes” collected from 1267 public human gut microbiome samples plus an additional 922 complete annotated prokaryotic genomes isolated and sequenced from human samples. About half of the genes contain KEGG function IDs in their annotation [14].

Bowtie2 [15] was used for alignment of MGS reads to a database of sequences from IGC, producing sequence matches at >90% identity at the default stringency (‐‐end-to-end ‐‐sensitive). DNA-DNA matches at this level of identity represent exact matches (DNA fragments from the same species) and orthologs between closely related species [16], so functional annotations can be confidently transferred from IGC/HOMD genes to a sequence reads in the MGS dataset.

The pipeline begins with a quality control check using FastQC [17], trimming of sequencing primers and low quality sequence with Trimmomatic [18], and removal of human sequences by Bowtie2 alignment to the human GRCh38 reference genome. All human genome sequence data filtered from the data files was discarded from this study. The trimmed non-human reads are then processed both by MetaPhlAn [19] to estimate the abundance of microbial taxa and by Bowtie2 alignment to the IGC/HOMD database to assign KEGG protein function IDs to each read (Figure 1). The MGS-Fast pipeline can process a typical MGS sample in about 4 hours on an 8 CPU Linux computer.

**Figure 1.**
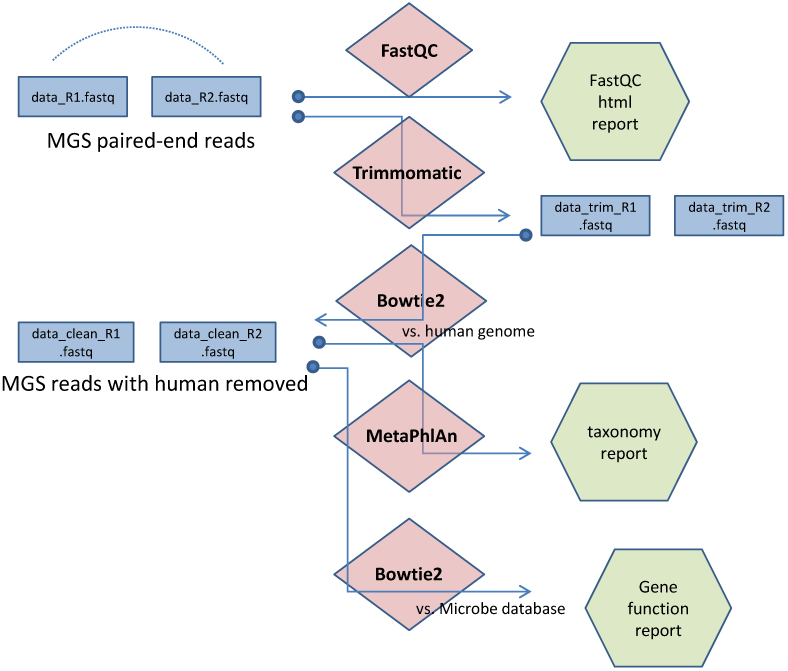
Flowchart representation of the MGS-Fast pipeline. Raw data as shotgun sequence FASTQ files is QC checked by FASTQC, then adapters and low quality sequences are removed by Trimmomatic. Human sequences are removed by Bowtie2 alignment to the human reference genome. The non-human reads are then processed by MetaPhaAn to estimate the abundance of microbial taxa and also by Bowtie2 alignment to the IGC/HOMD database to assign KEGG protein function IDs.

The MGS-Fast metagenomics data analysis pipeline has been implemented within a Docker virtual machine (VM) container [20], which has been made publicly available for installation on a variety of computing environments. With this approach we provide easy access to a pre-configured version of the data analysis pipeline (Suppl. 1), that does not required any installation other than downloading the container (and reference data sets). The Docker VM can be run on a local computer, shared compute cluster, or on-demand cloud computing platforms to scale data analysis. Our VM also provides a built-in Galaxy server as a graphical web-based user interface to run the pipeline and managing the data inputs and outputs. By starting the container and accessing the Galaxy interface (Figure 2 and Suppl. 1), users can access individual bioinformatic tools or the complete MGS-Fast data pipeline, as well as simple tools to import data and visualize analysis results.

**Figure 2.**
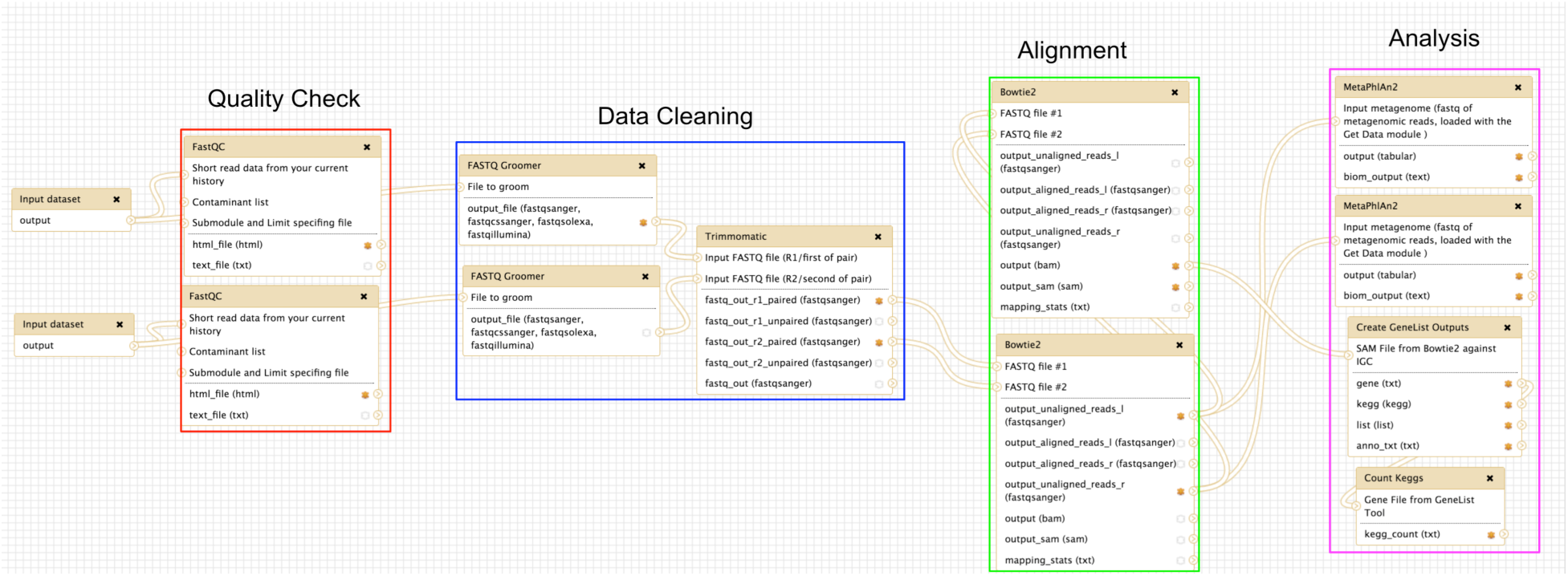
The MGS-FAST pipeline implemented as a Galaxy workflow.

## Results

We evaluated Bowtie2 alignment of various metagenomics data sets and controls to the IGC/HOMD database by the percentage of reads aligned (Table 1). A high percentage of aligned reads results in annotation of many reads. Low alignment percentage leaves many reads unannotated, but is also evidence of low false positives for a dataset that does not contain sequences from human microbiome organisms. The MGS-Fast pipeline was developed for the analysis of human upper GI tract samples, where an average of 49% (SD 5.7) of FASTQ reads map to the IGC database. Human gut (fecal) samples map to IGC at 95%, but human oral microbiome samples map at 38% and human skin at 35%. Interestingly, mouse fecal samples map at 60%, so the difference in local environment has more effect on the microbial community composition than host species. False positive matches were evaluated by aligning a set of randomly generated DNA sequences as a FASTQ file generated by the XS simulator, which had only 0.5% alignment. As a positive control, we mapped simulated reads from the *E. coli* K12 reference genome (GenBank: accession U00096.3) which aligned at 98.5%. Simulated FASTQ reads from the human reference genome GRCh38 aligned at 7.3%, which was somewhat surprising, since the construction of the IGC/HOMD database included a step to filter out human sequences. A metagenomic sample from copper mine waste (MG-RAST accession 4664533.3) aligned at only 8.69%, and a synthetic metagenome (SRR3732372) made from a mixture of DNA from lab strains of bacteria aligned at 9.75%. The HMP mock community (SRR172902) mapped at 28.8%.

**Table 1.** Alignment of FASTQ reads to the IGC catalog with Bowtie2.

| Source of Sample | Sample ID | Percent Alignment to IGC with Bowtie2 |
| --- | --- | --- |
| Human gut metagenome | SRR2822459 | 95.62 |
| Mouse gut metagenome | MG-RAST 4535626.3 | 89.71 |
| Human oral metagenome | SRR769511 | 37.97 |
| Human esophagus metagenome | (dbGaP phs000260.v3.p2) | 49.36% (SD 5.7) |
| Human skin metagenome | SRR1646957 | 35.81 |
| Human genome GRCh38 | FASTQ simulated by MetaSim | 7.35<br>(an estimate of false positives) |
| E coli K12 genome | FASTQ simulated by MetaSim | 98.5 |
| Synthetic microbial community | SRR3732372 | 9.75 |
| HMP Mock Community | SRR172902 | 28.82 |
| Copper Mine waste | MG-RAST 4664533.3 | 8.69 |
| Random simulated genomic sequence | generated by XS simulator | 0.53 |

The MGS-Fast pipeline was applied to 10 liver cirrhosis and 10 control samples from the study by Qin et al [8] (Figure 3). MGS reads were aligned to IGC/HOMD genes and assigned the corresponding KEGG IDs, producing gene function abundance counts for each sample. Following the recommendations of McMurdie and Holmes [21], this data was analyzed as a mixture model with a Negative Binomial distribution, so the between group differences of KEGG ID abundances can be calculated with edgeR software [22]. A total of 502 KEGG IDs are significantly different (FDR corrected p-values > 0.05) between 10 liver cirrhosis and 10 control samples, (KEGG abundance scores, fold change and FDR corrected p-values are shown in Suppl. Table 2).

**Figure 3.**
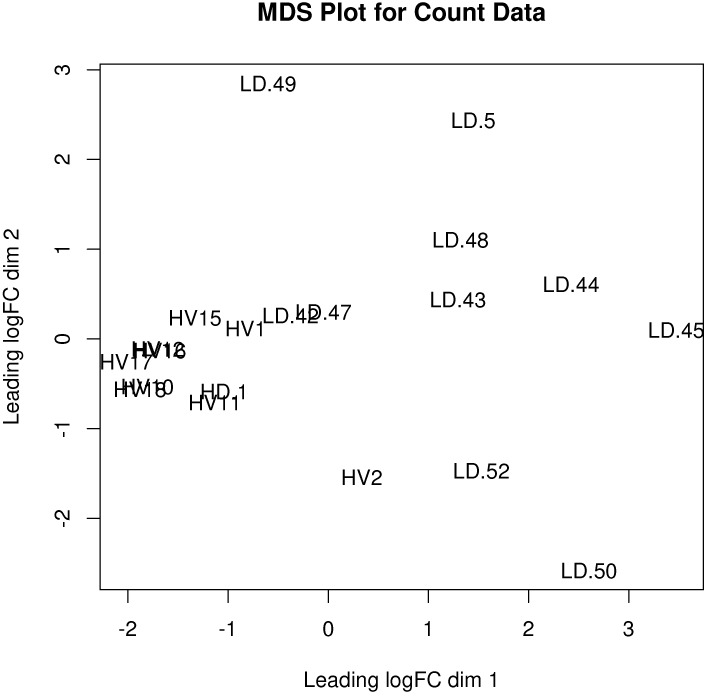
Diseased vs. Healthy Liver KEGG Abundance Plot.

## Discussion

Our goal for this method is to process MGS samples with a light computational load, and produce reliable functional mappings for metagenomic DNA sequence reads. Carr and Borenstein [23] compared MGS annotation using BLAST vs. BWA (a DNA sequence similarity tool very similar to Bowtie) and they conclude that at short evolutionary distances, BWA has a higher precision and recall than BLAST for identifying KEGG orthologs, but recall and precision for BWA drops dramatically at greater evolutionary distances. Bowtie2 is stringent in finding matches between reads and a target DNA database, requiring about 90% DNA sequence identity, but it is less sensitive than translated BLAST, which can utilize information from conservative amino acid substitutions. It is possible to increase the sensitivity of our method by changing the Bowtie2 parameters, such as “‐‐*very-sensitive-local*” or increasing the number of allowed mismatches. However, less stringent DNA-DNA alignments will create more false positives, and will make the assignment of metabolic function to MGS reads less reliable.

With our Docker virtual machine, we provide a pre-configured and ready-to-execute sequencing informatics solution that can be installable, replicated, and reused across computing platforms of all scales from institution-wide compute clusters and clouds to small data centers, or even laboratories that have only personal desktop computers.

Annotation of MGS reads with Bowtie2 requires a comprehensive database of annotated bacterial DNA sequences that are closely related to the species present in the sample. We provide a FTP/http server with a copy of the IGC database for human gut microbes in both FASTA and Bowtie2 index format (http://www.hpc.med.nyu.edu/~browns02/meta/). We have available a complete package that includes the hg38 in FASTA and Bowtie2 index format, IGC in FASTA and Bowtie2 format and sample datasets (http://bioitcore.hunter.cuny.edu:9988/metagenomics_package.tar.gz). Investigators that are working with samples very different from human gut can create their own custom database of genes by *de novo* assembly of multiple individual shotgun data sets and annotation of the genes found in these assemblies. Then a custom database of microbial genes can serve as a bridge between the FASTQ reads from each sample and the gene functions, increasing accuracy and reducing compute time for functional assignment of reads in new samples. A public reference collection of microbiome gene catalogs for many human body sites could be used to greatly accelerate microbial functional analysis by metagenomic shotgun sequencing.

## Abbreviations

KEGG: Kyoto Encyclopedia of Genes and Genomes
MGS: Metagenomics Shotgun sequencing
HMP: Human Microbiome Project
IGC: Integrated Gene Catalog of the human gut microbiome

## Supplemental Files

Supplement 1. The MGS-Fast Docker pipeline, installation and use instructions.

Supplement 2. Table of differential KEGG ID abundances for liver cirrhosis vs control shotgun sequencing samples.

## Declarations

Availability of data and material: The MGS-Fast software pipeline is available as a Docker image from docker pull bcil/metagenome:nyu_3.0 > /dev/null. Detailed instructions on the use of the Docker system and installation and use of the MGS-Fast image are available in Supplement 1 of this manuscript.

Data used in our evaluation studies are available from NCBI SRA (SRR1646957, SRR769511, SRR2822459, SRR3732372, SRR172902) and MG-RAST accessions (4535626.3, 4664533.3) as described in the text. The reference genomes for human GRCh38.p9 and *E. coli* K12 are available from GenBank (GCA_000001405.24, U00096.3)

Data from the HMP Foregut microbiome study are available from NCBI dbGAP phs000260.v3.p2 under Authorized Access restrictions.

## Author Contributions

The construction and testing of the MGS-Fast method was conducted by S. Brown with assistance from Y. Hao and H. Chen. The Docker image for MGS-Fast was built by B. Laungani with assistance from K. Krampis, T. Ali, C. Dong, C. Lijeron, and B. Kim. The manuscript text was written by S. Brown. Supplement 1 was written jointly by B. Laungani with additions by K. Krampis, T. Ali, C. Dong, C. Lijeron, and B. Kim.

## Acknowledgements

XXX

